# Deformable image registration for automatic muscle segmentation and the generation of augmented imaging datasets

**DOI:** 10.1101/2022.08.09.503405

**Authors:** William H. Henson, Claudia Mazzá, Enrico Dall’Ara

## Abstract

Muscle segmentation is a process relied upon to gather medical image-based muscle characterisation, useful in directly assessing muscle volume and geometry, that can be used as inputs to musculoskeletal modelling pipelines. Manual or semi-automatic techniques are typically employed to segment the muscles and quantify their properties, but they require significant manual labour and incur operator repeatability issues. In this study an automatic process is presented, aiming to segment all lower limb muscles from Magnetic Resonance (MR) imaging data simultaneously using three-dimensional (3D) deformable image registration. Twenty-three of the major lower limb skeletal muscles were segmented from five subjects, with an average Dice similarity coefficient of 0.72, and average absolute relative volume error of 12.7% (average relative volume error of -2.2%) considering the optimal subject combinations. Segmented MR imaging datasets of the lower limb are not widely available in the literature, limiting the potential of new, probabilistic methods such as deep learning to be used in the context of muscle segmentation. In this work, Non-linear deformable image registration is used to generate 69 manually checked, segmented, 3D, artificial datasets, allowing access for future studies to use these new methods, with a large amount of reliable reference data.

## 1. Introduction

Muscles enable all elected movements of the human body [1]. Relationships between structural muscle characteristics such as muscle volume, geometry and length, or level of fatty infiltration and the functional capacity of individual muscles have long been established [2,3,4]. Specifically, muscle volume and geometry are indicative of the maximal force that a muscle is capable of outputting [3,5,6] and fat infiltration within muscle tissue, known as myosteosis, reduces the saturation of contractile tissue, hindering the force generating capacity of a muscle [3,4]. Longitudinal changes in these structural characteristics are recognised as components of both aging [7,8,9] and the development of musculoskeletal (MSK) and neuromusculoskeletal disorders [10,11,12,13,14]. Through medical imaging analysis, structural muscle characteristics are measurable *in vivo* in a process named muscle segmentation [10,11].

Both Computed Tomography (CT) and Magnetic Resonance (MR) imaging, have been used to non-invasively gather quantitative structural muscle characteristics such as volume [15,16] or geometric shape [15,16,17]. The structural characteristics of the skeletal muscles within the lower limb are of particular interest, due to their capacity to enable locomotion [1,15,19]. As the lower limbs are such a large area of the body, many studies prefer MR imaging over CT, to limit ionising radiation exposure of subjects enrolled in studies or of patients in future potential clinical applications [15, 19]. The current approach used within the literature to gather these structural muscle characteristics from MR images is manual segmentation, during which the operator defines in each slice of the MR image (or in a subgroup of them) the contour of each muscle [15,19,20]. There are two main limitations of manual segmentation: the required operator input time and operator dependency issues of the outputs [5,6,15,18]. It is generally accepted that there are 35 individual lower limb muscles, not including those in the feet (some of these muscles can be separated into different branches or sub-muscles [15,22,23]), which must be segmented from (in the order of) hundreds of images, incurring a high processing time [15,20,21]. Recent advancements in computer vision (interpolation between segmented slices) and hardware (trackpads) have brought operator interaction time down to approximately 10 hours to segment all muscles within one lower limb [15,20,21]. Not only is this interaction time excessive, but operators must undergo training to achieve repeatable segmentation results from an intra-operator standpoint (± 10% volume is typically acceptable) [15]. Regardless of training, as suggested, there are significant inter-operator dependency issues noted within the literature, which have been shown to misinterpret muscle volume by up to 50% (for example, the peroneus brevis and longus [15]), depending on the muscle of interest and study cohort [15,21,24]. These limitations of manual segmentation prevent the utilisation of muscle segmentation as a technique to inform large-scale quantitative investigations into muscle characteristics.

Many different automatic segmentation methods have been investigated within the literature in recent years to replace the manual approach [21,24,25,27]. Statistical shape modelling (SSM) entails the generation of an average atlas geometry, which can be scaled or deformed to fit individuals. It has been used to achieve a good agreement of automatically and manually generated segmentations of single muscles from MR images, such as the quadratus lumborum within the lower back by Engstrom et al. [25] where the Dice similarity coefficient (DSC), a volumetric and spatial measure of agreement, achieved was 0.86 ± 0.08 (mean ± standard deviation). This technique was shown to be well suited to the automatic segmentation of this muscle given its non-complex, truncated cone-like shape, but has not been explored in the segmentation of many individual muscles simultaneously. The large variability of muscle volume and geometry within the lower limb skeletal muscles between subjects, even within cohorts with similar anthropometric characteristics, limits the application of SSM to segment these muscles [15,29,30,31]. Image registration has also been explored within the literature to perform muscle segmentation. Simplistic applications, such as two-dimensional (2D) deformable image registration between subsequent MR imaging slices within subjects has been used to propagate segmentations of individual slices into partial sections of 3D muscle geometry using only a few manually segmented slices, with encouraging results (DSC ≈ 0.91) [32]. 3D image registration has been used within longitudinal studies to populate MR images with partial segmentations of a small number of muscles to good effect, such as within the studies presented by Le Troter et al. [33] and Fontana et al. [34]. Though this longitudinal approach provides insight into the change in muscle characteristics over time, multiple MR image sequences are required from individual subjects at two different timepoints and one dataset must be manually segmented. Within the literature, inter-subject registration aiming to segment the muscles within a new subject, referencing a previously segmented subject has not yet been fully explored to the best of the author’s knowledge. An in-house image registration algorithm (Sheffield Image Registration Toolkit, ShIRT) has been used to segment both hard [43] and vascular [42,43] tissues with a high level of accuracy but has not yet been tested in the application of muscle segmentation. ShIRT performs deformable (non-linear) image registration, allowing high degrees of anatomical variability between inputted images to be addressed [41,42,43], and has the potential to automatically segment blocks of muscles or individual muscles given a fully segmented reference subject, but this is yet to be explored within the literature.

Other methods have been shown to be effective in the segmentation of muscles. Probabilistic machine learning methods such as deep learning have been used to automatically segment the 3D geometry of individual muscles from MR images taken from several different cohorts [20,21,24]. These methods employ Convolutional Neural Networks (CNNs) which learn patterns that identify important features from training data in order to apply these learned patterns to segment new, unseen data [20,21,24]. Notably, the methods recently proposed by Ni et al. [24], where all lower limb muscles within a cohort (n = 64) of young healthy athletes were segmented with DSC comparable to that of the inter-operator dependence (DSC ≈ 0.9), and those proposed by Zhu et al. [21] where all muscles within the shank were segmented from a cohort (n = 20) of children with cerebral palsy (DSC ≈ 0.88). Though the segmentation accuracy found within these studies is remarkable, these methods are not widely accessible due to the main limitation of current deep learning methods: the requirement of large training databases (minimum ∼20 segmented 3D images, the greater this number the more robust the method) [35]. Unfortunately, generating these segmented imaging datasets might not be well suited to MR imaging, given the associated high costs and manual processing time. Additionally, when used in medical image segmentation, deep learning generally has the major limitation of a significantly reduced performance when assessing imaging data taken from widely varying cohorts [35]. Data augmentation is a technique widely used in association with CNNs for the purpose of supplying greater amounts of training data and helping to generalise their application to image classification and segmentation tasks [36,37]. Within this context, image registration has been previously used to generate augmented images to facilitate the analysis of brain tumours [38] and skeletal deformities [39]. This suggests that, while not attempted before, similar approaches might be adopted for muscle segmentation.

The aim of this study is hence twofold. The first is to evaluate the accuracy of a method for automatic segmentation of individual skeletal muscles in the lower limb from MR imaging data using 3D deformable image registration. Secondly, the effectiveness of this approach in the generation of augmented datasets is explored.

## 2. Methods

### 2.1. Subjects & imaging acquisition method

Retrospectively available lower limb T1-weighted MR images from 11 post-menopausal women (mean (standard deviation): 69 (7) years old, 66.9 (7.7) kg, 159 (3) cm) were used for this study [15]. Images were collected using a Magnetom Avanto 1.5T scanner (Siemens, Erlangen Germany), with an echo time of 2.59 ms, repetition time of 7.64 ms, flip angle of 10 degrees. The study was approved by the East of England – Cambridgeshire and Hertfordshire Research Ethics Committee and the Health Research Authority (16/EE/0049). The MR images were acquired in four sequences, capturing the hip, thigh, knee, and shank. To reduce scanning time while still providing detailed geometries of the joints for use within the original study, the joints were acquired with a higher resolution (pixel size 1.05 mm^2^, slice spacing 3.00 mm) than the long bone sections (pixel size 1.15 mm^2^, slice spacing 5.00 mm). The sequences were stacked in MATLAB forming one continuous 3D image from hip to ankle, firstly by homogenising the resolution of each of the imaging sequences taken from the different sections to be 1.00×1.00×1.00 mm^3^ through tri-linear interpolation (interp3, MATLAB 2006a). The fields of view of the images across the four sequences were equated by wrapping the images in blank data (greyscale value of 0), referencing the spatial metadata of the images to retain the relative subject position across the imaging sequences for each subject. The homogenised sequences were concatenated in the longitudinal direction, removing half of any overlapping volume from each section where the fields of view overlapped. Lastly, the images were cut in half in the frontal axis, isolating only the right limb. A sub-cohort of 5 of the 11 subjects was selected for automatic segmentation. The five subjects were chosen with the aim of creating a sub-cohort with a wide anatomical diversity, choosing the tallest and shortest [154.0 cm, 164.2 cm], the subjects with the lowest and highest Body Mass Index (BMI, kg/m^2^) [21.2, 32.1], and the youngest and oldest participants [59, 83]. Each subject was used as both a target and a reference for the image registration algorithm, creating 20 subject pairings for the sub-cohort (inter-subject analysis). For comparison, the 5 subjects were registered with a procedure similar to the inter-subject analysis, using the opposing limb (left vs right) as the reference dataset for the registration (intra-subject analysis).

### 2.2. Reference segmentations

Each of the five subjects involved in this study were segmented manually, as presented by Montefiori et al. [15]. Within this database, the muscles for which the coefficient of variation of the manual segmentations when repeated by the same operator on three separate runs was greater than 10% were removed from the study, reducing the number of muscles considered in this study from 35 to 23. Table 1 presents the range of volumes of the 23 muscles considered within this study within the cohort of 5 subjects. Their manual muscle segmentations were used as the templates to populate imaging data of new subjects with automatically generated muscle segmentations through image registration and to validate them.

**Table 1:**
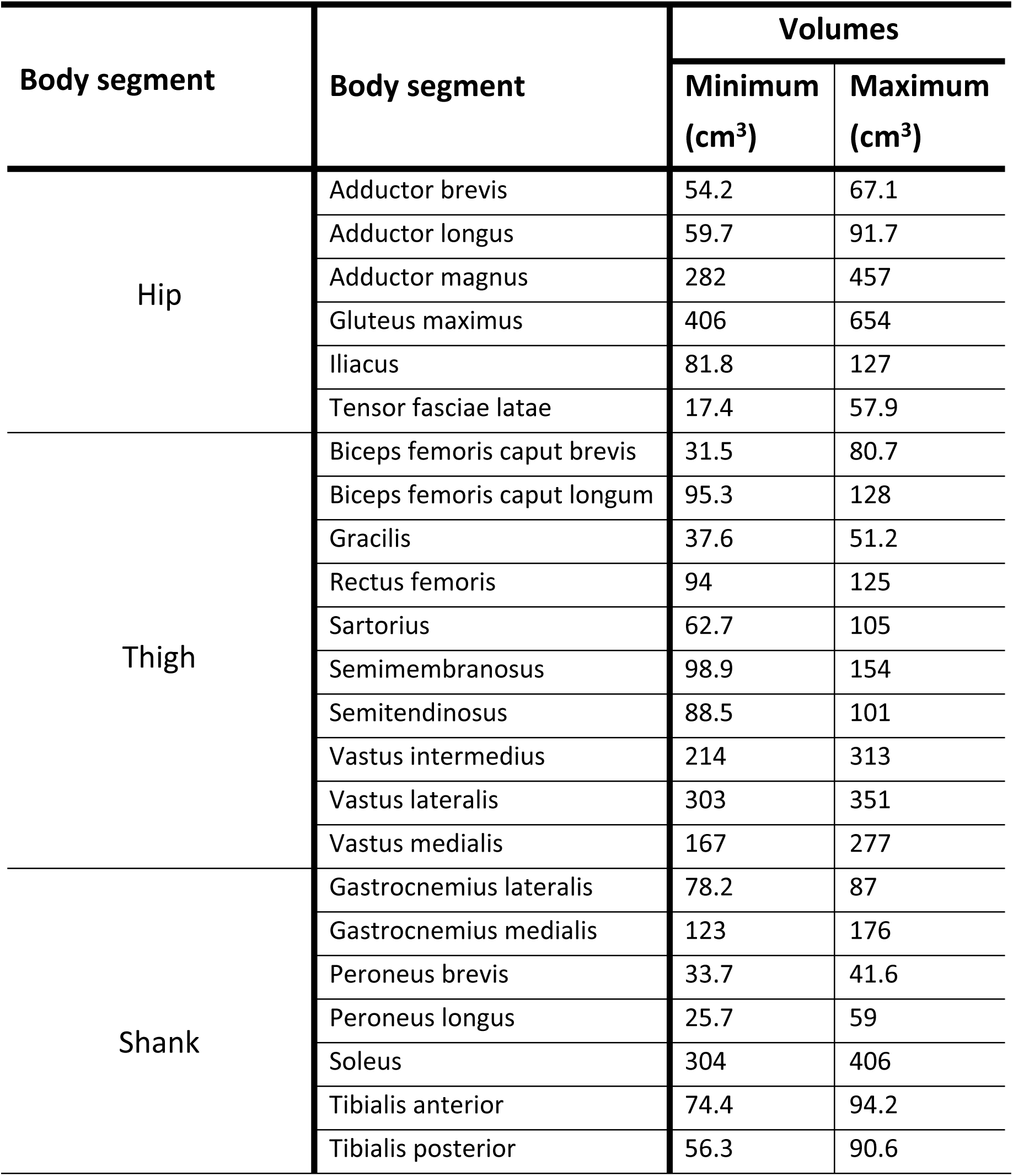
The range of volumes of the muscles included within the study for the 5 subjects considered. The muscles are separated into three sections of the body (hip, thigh, and shank). The muscles considered are those that were segmented with an acceptable level of repeatability [15]. Full description of muscle volumes within each subject expanded upon in supplementary material 1.

| Body segment | Body segment | Volumes |  |
| --- | --- | --- | --- |
|  |  | Minimum (cm<sup>3</sup>) | Maximum (cm<sup>3</sup>) |
| Hip | Adductor brevis | 54.2 | 67.1 |
|  | Adductor longus | 59.7 | 91.7 |
|  | Adductor magnus | 282 | 457 |
|  | Gluteus maximus | 406 | 654 |
|  | Iliacus | 81.8 | 127 |
|  | Tensor fasciae latae | 17.4 | 57.9 |
| Thigh | Biceps femoris caput brevis | 31.5 | 80.7 |
|  | Biceps femoris caput longum | 95.3 | 128 |
|  | Gracilis | 37.6 | 51.2 |
|  | Rectus femoris | 94 | 125 |
|  | Sartorius | 62.7 | 105 |
|  | Semimembranosus | 98.9 | 154 |
|  | Semitendinosus | 88.5 | 101 |
|  | Vastus intermedius | 214 | 313 |
|  | Vastus lateralis | 303 | 351 |
|  | Vastus medialis | 167 | 277 |
| Shank | Gastrocnemius lateralis | 78.2 | 87 |
|  | Gastrocnemius medialis | 123 | 176 |
|  | Peroneus brevis | 33.7 | 41.6 |
|  | Peroneus longus | 25.7 | 59 |
|  | Soleus | 304 | 406 |
|  | Tibialis anterior | 74.4 | 94.2 |
|  | Tibialis posterior | 56.3 | 90.6 |

### 2.3. Image pre-processing

The MR images of each subject were pre-processed to homogenise the distribution of fat tissue within the scans and maintain the focus of registration to the muscles. For each 2D slice of imaging data (example slice shown in Fig. 1. A) within each subject, firstly, the air-skin boundary was located using a Canny edge detector [40]. The area within the skin boundary was filtered (Fig. 1. C), in response to a threshold established from the greyscale frequency intensity plots of the images, creating a mask that contained only the muscle tissue (Fig. 1.D). A layer of fat was wrapped around the muscle tissue (Fig. 1.E and 1.F) to emphasise the outer boundary of the muscle tissue. The depth of this layer of fat was made equal to the optimal nodal spacing (NS, a parameter of the registration [41], set to 5 mm, details in 2.4) as the registration operates optimally in the circumstance that the object being registered is of similar size to the NS [41]. There were two possible scenarios for the fat wrapping process: 1) the layer of fat within the image was greater than 5 mm, and 2) the layer of fat was less than 5 mm. In the first scenario, the subject’s fat tissue was wrapped around the muscle tissue at a depth of 5 mm. In the second scenario, artificial fat was wrapped around the body which was built in response to the greyscale frequency intensity peak that represents the fat. The pixels within 5 mm of the muscle tissue that lay outside the body were randomly assigned values using a uniform distribution with minimum and maximum equal to the mean ± standard deviation of the frequency intensity peak representing the fat.

**Fig. 1:**
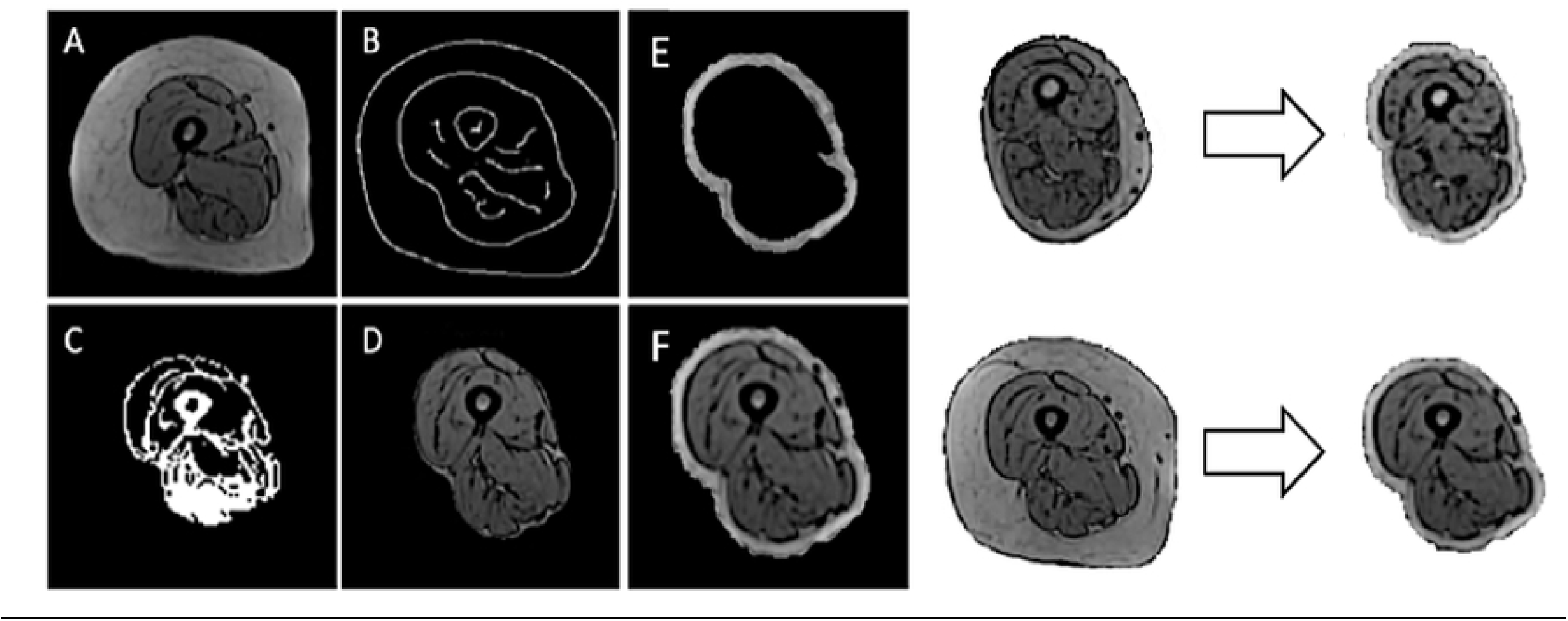
The process of masking the fat tissue surrounding the muscles from the raw MR images (left) and wrapping in a homogenous layer of fat for two images taken from different subjects (right). The subject along the top row (right) had a fat layer less that 5 mm thick and was wrapped with artificial fat, where the subject along the bottom row had a depth that was sufficient.

### 2.4. Segmentation

Following pre-processing, subject imaging datasets were registered using ShIRT [41]. In the registration process, displacement functions were computed that map each pixel in a reference image to a corresponding pixel in the target image. ShIRT solves displacement equations at nodes of an isotropic hexahedral grid overlapped to the fixed and moved images, with distance between the nodes equal to NS. The optimal NS for this registration task was found through a sensitivity analysis (see supplementary material 2). Throughout the registration process the optimal nodal displacements are smoothed in response to a smoothing coefficient, optimised in each registration to solve the registration problem [41] (this was verified to be indeed optimal for this application, see sensitivity analysis in supplementary material 2). The 3D displacement field is calculated using tri-linear interpolated displacements between the nodes of the grid. The registered image was generated after applying the transformation to the reference image and using tri-linear interpolation. Similarly, the automatic segmentation of the muscles within the target subject was calculated applying the transformation to the manual segmentations of the reference subject (Fig. 2).

**Fig. 2:**
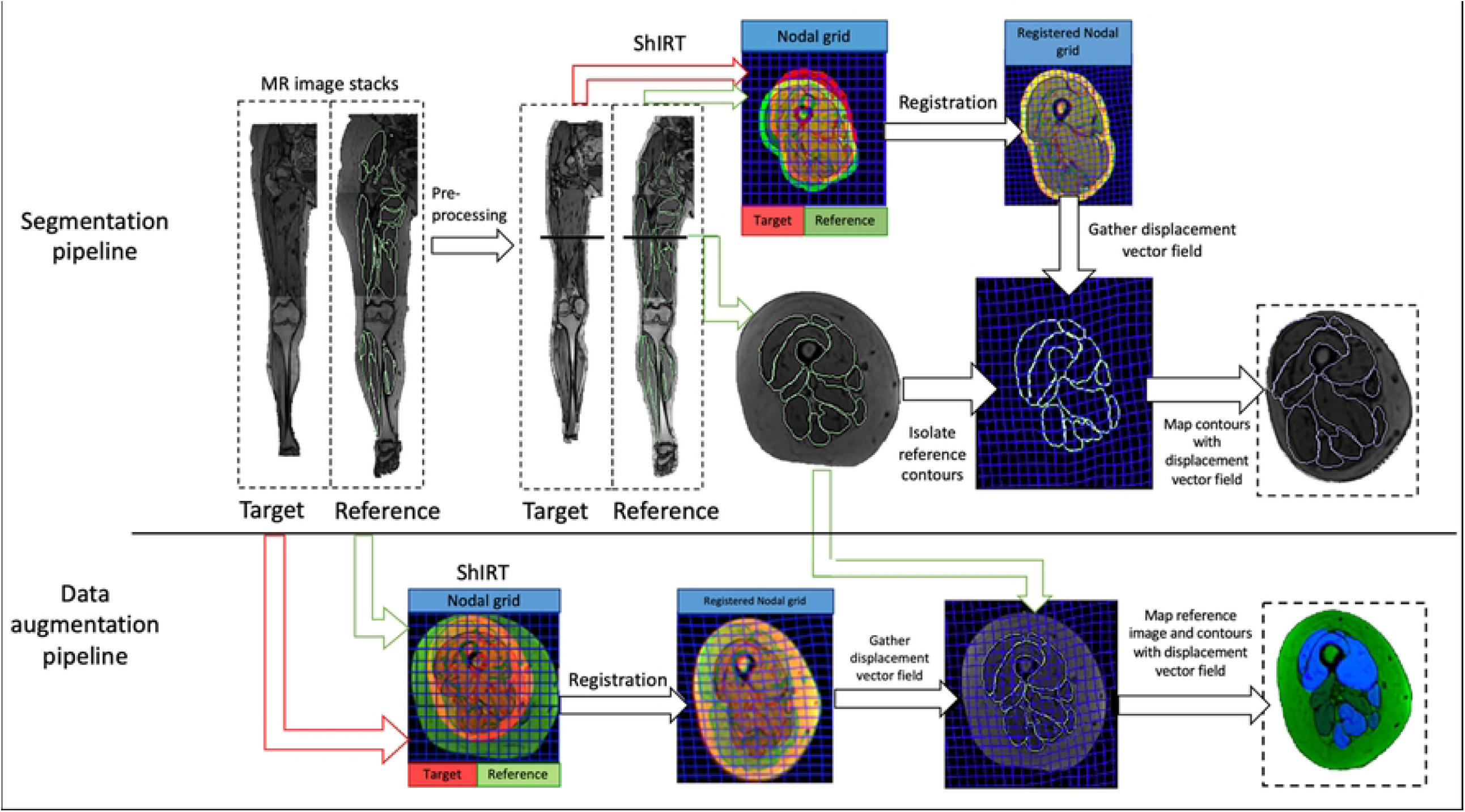
The image registration process, shown for one 2D slice of imaging data (location within imaging sequences highlighted with a black line). Segmentation pipeline: the target and reference subject were pre-processed, homogenizing the fat layer, and registered in ShIRT. The map found through registration was applied to the manual segmentation contours of the reference subject (shown in green), resulting in an automatic segmentation of the target subject (shown in blue). Data augmentation pipeline: The combined MR imaging sequences are registered in ShIRT. The map outputted from the registration was used to deform the reference subject’s 3D imaging data and reference manual segmentations, resulting in a fully segmented, augmented 3D image. The augmented images are shown with each muscle taking a different greyscale value (visualised in blue image channel).

To gauge the accuracy of the resulting segmentations, the registration and segmentation pipeline was used to segment the right limbs of the 5 subjects using the opposing limb as the reference input. The muscles within opposing limbs have been proven to be anatomically similar but distinct in both volume and geometry [15]. For these reasons, using the opposing limb in the segmentation pipeline should provide the best possible reference for the segmentation of the muscles within each of the 5 subjects.

### 2.5. Segmentation validation

The reference registered image and the target image were overlapped to assess the quality of the registration. The two images were visualised simultaneously, with the registered and target images shown in green and red, respectively. Well registered images appear yellow with very few green or red flecks. Fig. 3 presents three example registration results, where the quality of registration increases from left to right.

**Fig. 3:**
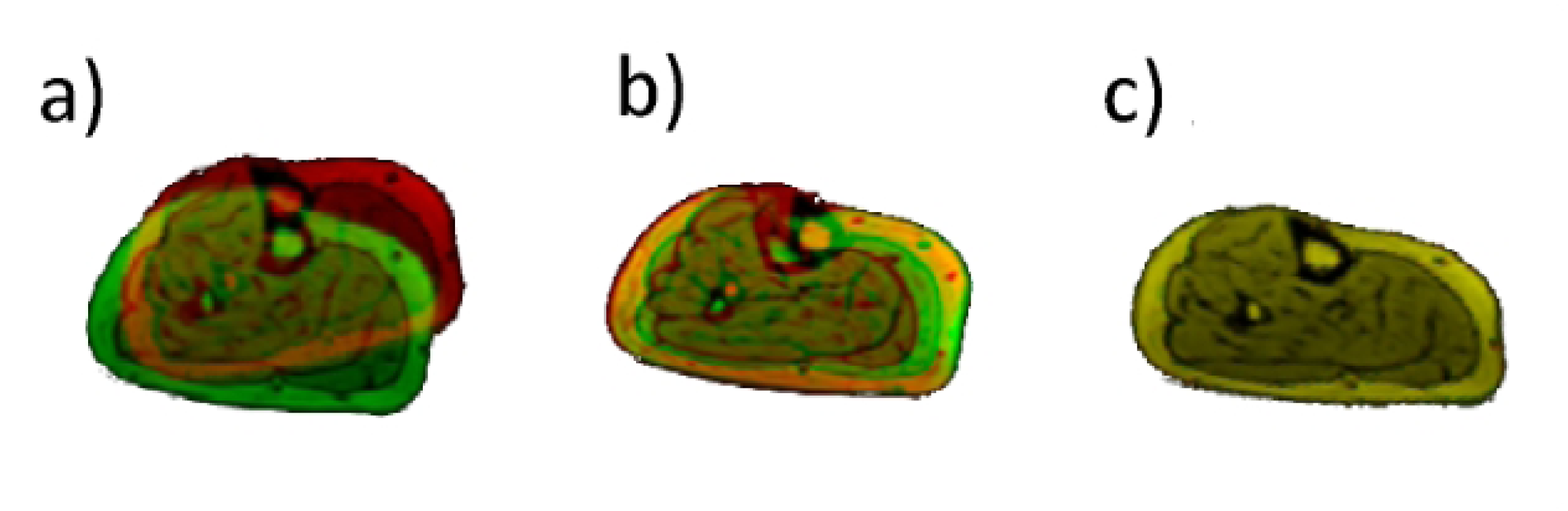
Registration results of images taken from the shank. The registration quality is visualised within these plots with poor, moderate, and flawless registrations shown in a, b, and c, respectively. Yellow colour represents well registered regions.

Three complementary quantitative metrics were used to test the accuracy of the automatic segmentation protocol. The relative volume error (RVE) was calculated following equation 1 for each muscle in each subject. Additionally, the total volume error (TVE) between the reference and automatically segmented muscles was calculated as the error between the sum of all muscle volumes, shown in equation 1.

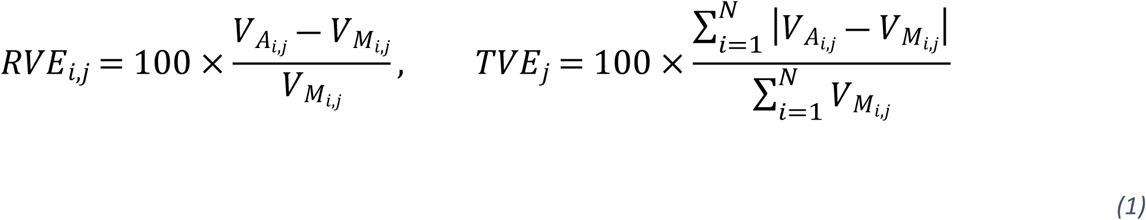

Where 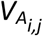 and 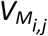 are the volumes of the automatic muscle segmentation and ground truth (manual) segmentations, respectively.

The Dice similarity coefficient (DSC) [45] was used to assess the accuracy of segmentation considering both volume and geometry, through comparison with the ground truth segmentation. The DSC varies between 0 and 1, with a value of 1 signifying that the proposed segmentation and ground truth are identical. The DSC was calculated (Equation 2) for each muscle (i) in each subject (j), where *A*_*i,j*_ and *M*_*i,j*_ represent the automatic muscle segmentation and the ground truth segmentation, respectively.

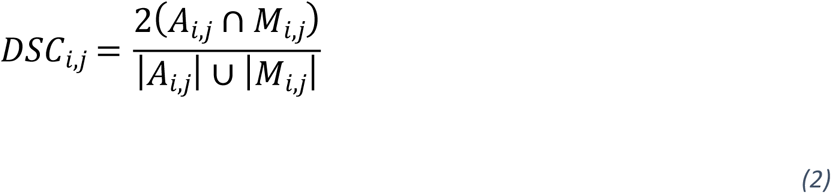

Finally, the Hausdorff distance (HD) [46] between the automatic and reference muscle segmentations was calculated for each muscle in each subject, following equation 3, where *a*_*i,j*_ is an element of *A*_*i,j*,_ *m*_*i,j*_, is an element of *M*_*i,j*_, and *d* is the magnitude of the minimum distance between *a*_*i,j*_ or *m*_*i,j*_ and the nearest neighbouring point within *M*_*i,j*_ or *A*_*i,j*_, respectively. For each subject the HD was calculated as the maximum among the minimum distances between the automatic and ground truth segmentations.

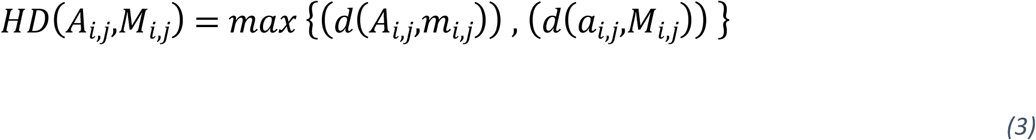

### 2.6. Generation of augmented data

The deformable image registration algorithm was used to generate segmented augmented MR imaging data, available for download within supplementary material 3 The stacked MR imaging data from the right limb of the 11 participants were registered to each of the other subjects in the cohort, giving 110 combinations. No pre-processing was applied. The displacement vector field outputted from ShIRT (Fig. 2) was used to deform both the MR imaging sequence and the manual muscle segmentations of the reference subject. The output of each of these processes was a fully segmented 3D image that was dissimilar to both the reference subject and the target subject (Fig. 2). A four-point criterion was used for checking both the images and the segmentations to ensure anatomical credibility of the augmented dataset: a) the boundaries of the long bones and the skin must be reasonably smooth and continuous; b) the positioning and orientation of the joints must be anatomically viable, with the bones fitting together realistically; c) the muscle segmentations should reflect the muscle structure; and d) the location of each of the muscles relative to one another must be realistic (e.g. the vastus lateralis must be lateral with respect to the vastus medialis). If any one of these criteria were not met, the augmented dataset was discarded. Out of the retained datasets, 15 chosen at random were retested by a different operator to confirm the specificity of the inclusion criteria. Finally, the available muscle volumes were compared from within the augmented and original databases. The mean volume within each database was computed for each of the 23 muscles considered. The difference between the volume of each muscle within the database and the average was then calculated, and this value was normalised against the mean volume. The resulting values were percentages representing the distribution of available muscle volumes within each database, which after normalisation, could be compared.

## 3. Results

### 3.1. Segmentation results

A visualisation of an example registration and of the results of one segmentation are highlighted in Fig. 4 for images taken from the hip, thigh, and shank, respectively. While the deformable image registration has accurately identified the muscle tissue in the target subject in most cases (yellow), some regions were not correctly registered (red or green). The segmentation results reflect this, where the registration appears successful overall, and the automatic segmentations are geometrically very similar to the reference segmentations. There are areas within the automatic segmentations that do not reflect the reference segmentations, such as the gluteus maximus in the hip section, and the tibialis muscles within the shank section. The automatic segmentations within the thigh section mostly agree with the reference segmentations.

**Fig. 4:**
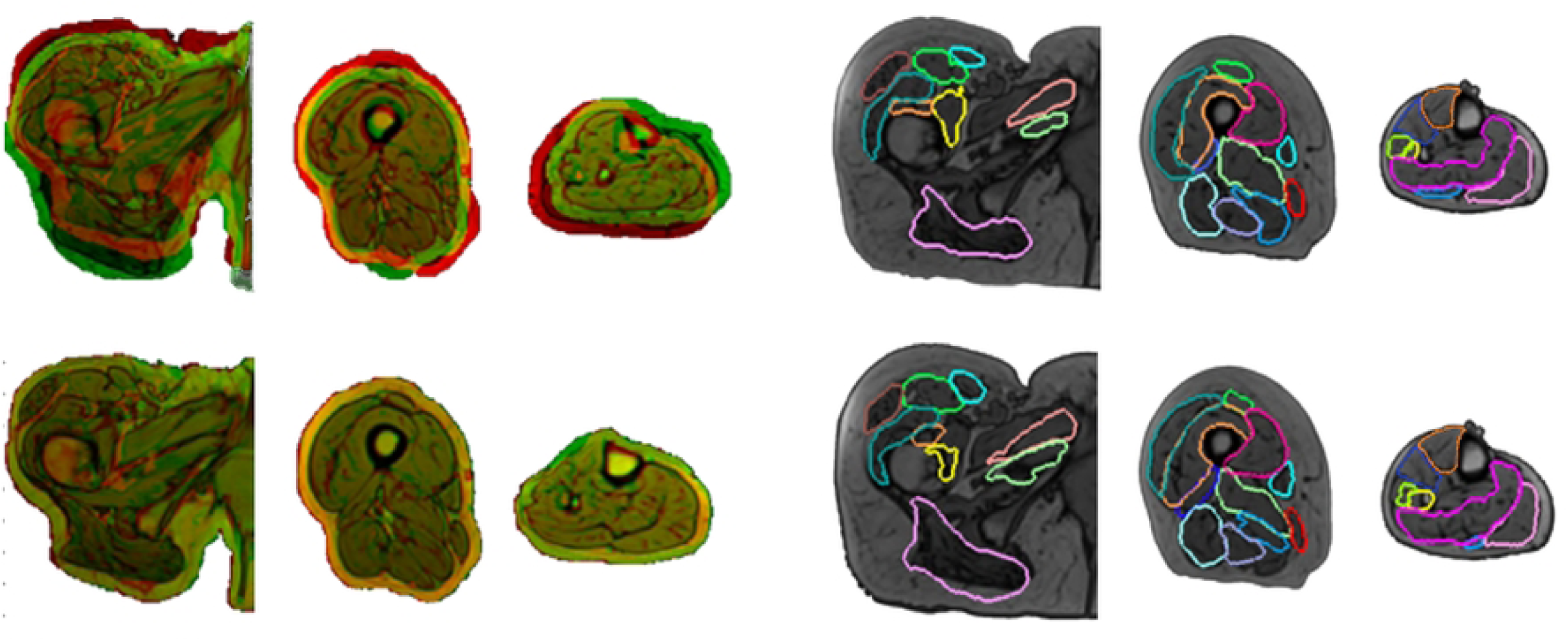
Registration and segmentation results from the combination of subjects resulting in the median average DSC (subject 4 and 2 as the target and reference, respectively). The registration inputs (top row) and outputs (bottom row) for these combinations of subjects are shown on the group of images on the left. The segmentation results are shown in the right three image groups, where the reference and automatic segmentations for the target subject are shown in blue and red respectively. The muscles that are not highlighted within the images, were found not to be segmented with an appropriate level of repeatability.

#### 3.1.1. Volume error

The TVE for the entire muscle body was 8.2 ± 5.1 % (mean ± standard deviation) across all subject combinations (Fig. 5). The mean RVE for the individual muscles was found to be below 12.8% for all combinations and all upper quartiles were below 40% error. The best performing combination was subject 5 with 1 as the target and reference respectively, with among the smallest mean (−2.2%) and with the lowest quartiles (lower and upper quartiles of -10.5% and 6.4%, respectively). The relative volume error was consistent across all muscles, with no correlation found between muscle volume and relative volume error (R^2^=0.092, p-value=0.159); the muscles with the highest variability within this cohort (adductor brevis, rectus femoris) made up the outliers within the distributions of RVE. The mean RVE from the intra-subject (left vs right) analysis was 0.35%, (Fig. 5).

**Fig. 1:**
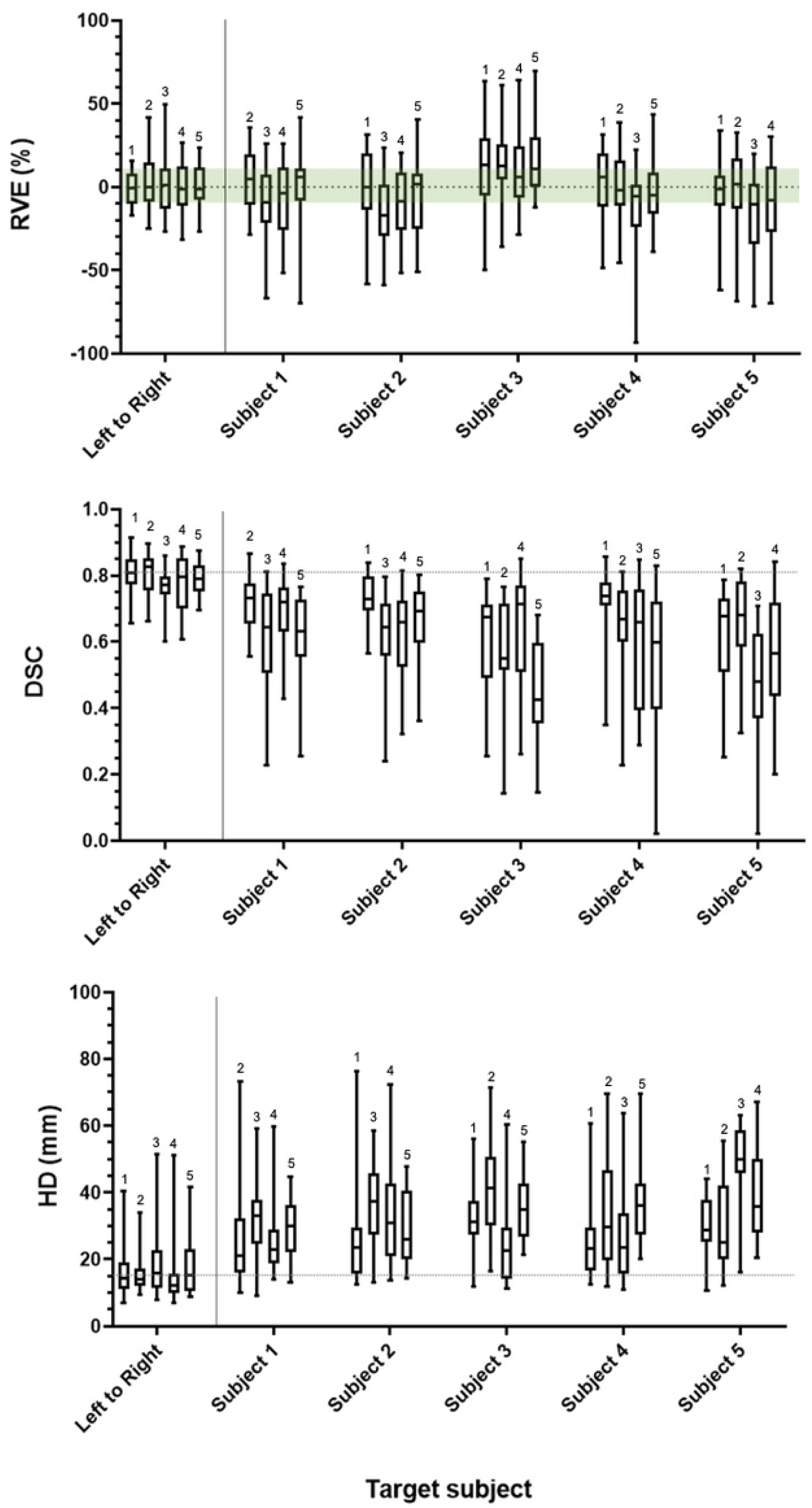
Relative volume error (%) (top), Dice similarity coefficient (centre), and Hausdorff distances (mm) (bottom) found for each muscle in each subject, using the other subjects in the sub-cohort as the reference. The numbers above each of the boxplots denotes the reference subject pair for each target subject 1-5 used within the registration. The green area represents the acceptable level of RVE resulting from inter-operator dependence, prescribed by Montefiori et al. 2019 [15]. The grey dashed lines represent the mean values from the intra-subject analysis for comparison. The box and whisker plots show the mean, interquartile ranges, and ranges across the 23 muscles considered.

#### 3.1.2. Dice Similarity Coefficient

When looking at the segmentations of the five subjects obtained using the other four as reference subjects, very variable results were observed. The greatest average DSCs were those resulting from the segmentation of subjects 1,2, and 4, using subject 2, 1 and 1 as the reference subject, respectively. The mean DSCs found for these combinations of subjects were greater than 0.70, lower quartiles greater than 0.67, and with a wide spread of results (0.35 < *DSC* < 0.88). Subjects 3 and 5 were segmented with a consistently lower DSC, with the average DSC considering all reference subjects found to be 0.61 and 0.60 respectively (0.69, 0.69 and 0.67 for subject 1, 2, and 4, respectively). Additionally, one of these subjects was always the worst performing reference subject considering DSC when used to segment all target subjects, with the lowest average DSC. There was a weak correlation found between muscle volume and the DSC of the automatic segmentations (R^2^=0.332, p-value=0.003). The average DSC found within the intra-subject analysis was 0.80 (Fig. 5).

#### 3.1.3. Hausdorff distance

Overall, the average HD was typically between 15 mm and 30 mm, with the upper quartile being below 40 mm, other than the segmentations of subject 3 and 5 using subject 2 and 3 as references, respectively (Fig. 5). The spread of results was large, with Interquartile Ranges (IQR) being between 7 mm and 21 mm. There was no correlation found between the HD and the size of the muscle for which the HD was calculated (R^2^=0.097, p-value=0.089), the error was consistent across muscles of all sizes. The average HD found within the intra-subject analysis was 17.7 mm.

### 3.2. Augmented data

After initial checking by the author, 69 of the 110 generated augmented datasets passed the inclusion criteria. 15 datasets were rechecked by an expert in muscle segmentation and all 15 passed, giving 100% specificity. Fig. 6 showcases some examples of the augmented images collected. Visually, the augmented images are well segmented, and are dissimilar to the reference subjects, particularly in the second row of images, where the relative fat depth of the moving subject (green) is retained, but the cross-sectional area of the thigh is equated to the fixed subject (red). The misalignment of the muscle tissue within the registered images, visible as concentrations of either red or green colouration, establish a difference in the muscle geometry within the registered and original data. The augmented subjects generated for 1 target subject (subject 1) are presented within supplementary material 3, for visual comparison.

**Fig. 6:**
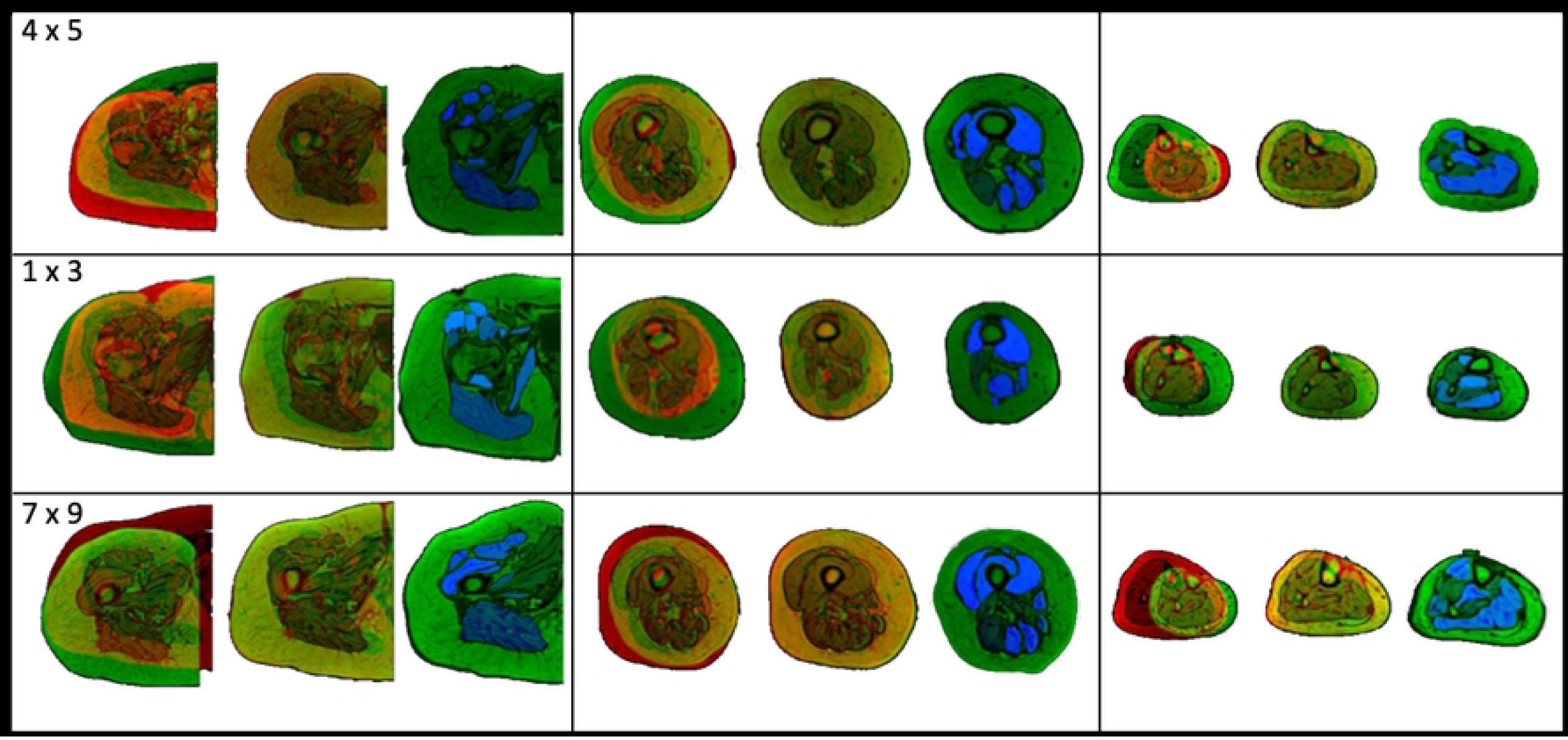
Inputs, outputs and resulting augmented subjects. Each row of images presents results within the hip (left), thigh (centre), and shank (right) for 3 subject combinations chosen at random (target x reference: 4 × 5 (top), 1 × 3 (middle), 7 × 9 (bottom)). Within each cell there are the inputted images into the registration (left), registered images with corresponding target image (centre) and resulting segmented, augmented images (right). The muscle labels are visible within the augmented images as the blue areas. Each muscle is assigned a distinct greyscale value and the labels are assigned alphabetically.

The anatomical variability of the muscles within the augmented database is compared to the original 11 subject database (Fig. 7). The volumes of each of the muscles within the original and augmented databases were normalised against the corresponding average muscle volume for each muscle within the respective databases. The percentage greater or smaller than the average volume was then calculated for each muscle, representing the variability of the muscle volumes within each database. The distributions of these percentages are presented (Fig. 7). The muscle volumes available within the augmented database were found to have a greater range of volumes, often 1.5 to 2 times greater than in the original database. The range of volumes for each muscle considered within the original and augmented databases are presented in supplementary material 4.

**Fig. 7:**
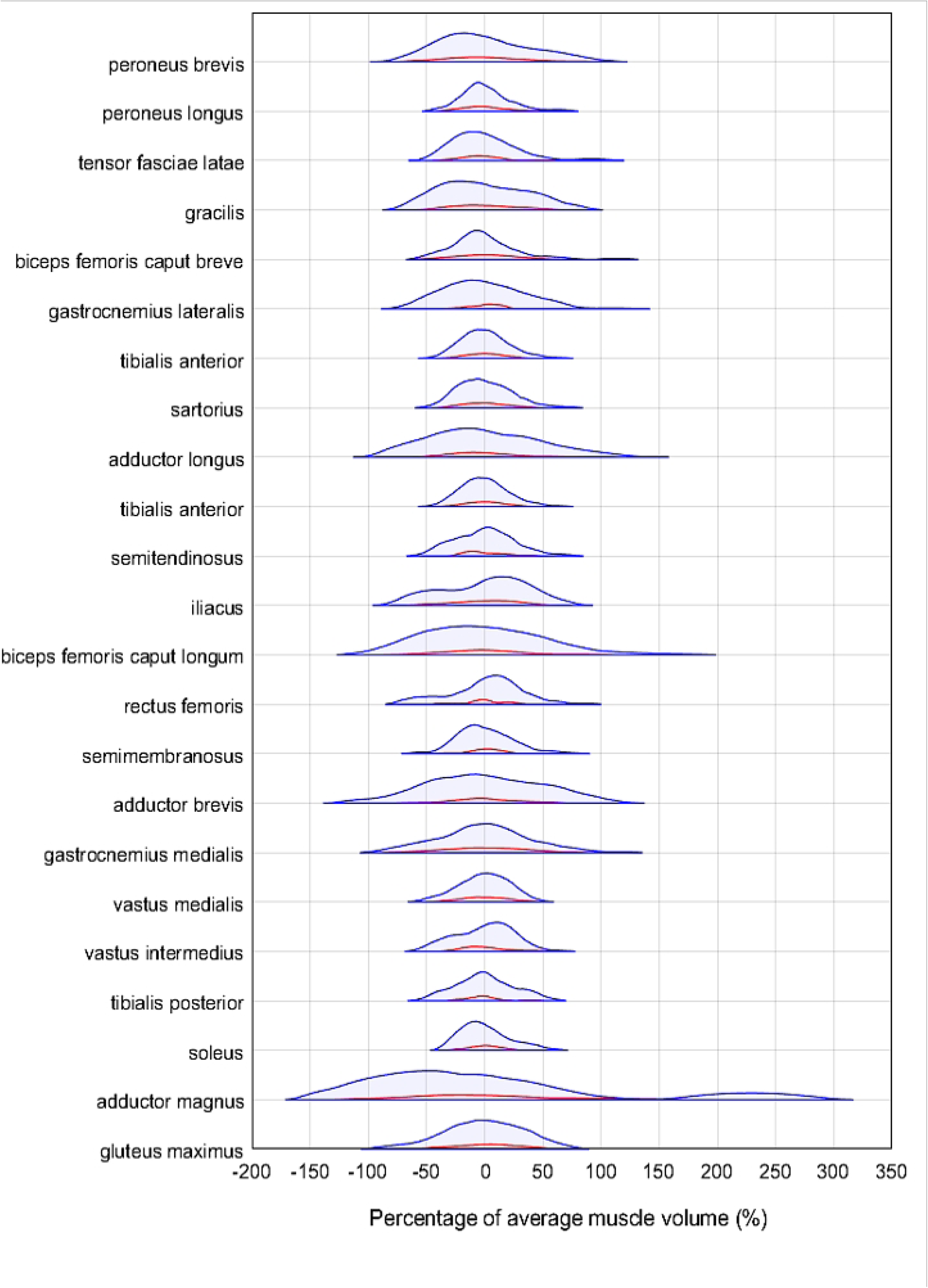
The anatomical variability of muscle volumes for each muscle, ordered from smallest to largest within the original and augmented databases shown in red and blue respectively. The height of the distributions was not normalised, and the violin plot contains 95% of the data, with 2.5% of data cut off from each side, removing outliers.

## 4. Discussion

This paper aimed at proposing a fully automatic tool to segment 23 major lower limb muscles simultaneously from MR imaging data using morphological image processing and deformable image registration. Furthermore, the same tool was used to generate a unique dataset including 69 fully segmented, augmented 3D images. To the best of the authors’ knowledge, this study represents the first attempt to segment complete 3D muscle geometry of many individual muscles simultaneously using deformable image registration while using different subjects as the reference. This would be desirable as muscle segmentation of a new subject could be performed without the need for manual processing.

All 23 muscles were segmented from five subjects with moderate success, considering three error metrics, the RVE, DSC and HD. The registration quality was high considering the combination of subjects that resulted in the median average DSC (Fig. 4) which suggests that in most cases, the registration performed as intended. This was confirmed by the total volume error metric, lower than 10% on average. However, all three-error metrics reflected a lower accuracy for the segmentation of individual muscles. The individual muscle RVE was typically larger than that of an acceptable level of inter-operator dependence (±10%) [15], with the lower and upper quartiles often exceeding ±10% in most subject combinations. The mean absolute RVE within the optimal subject combinations was 12.7%, meaning that on average, there was an over or underestimation of the muscle volume greater than the effect of operator variability. This indicates that the method would be best suited when only interested in the volume of the overall muscle body. Capturing the total muscle volume has proven useful in studies such as Handsfield et al. [19], where regression equations were presented, to estimate individual muscle volume from total muscle volume and other anthropometric data such as height and BMI. The DSC results, on the other end, indicate that if the purpose of the segmentation was that of extracting internal muscle characteristics, such as the level of fat infiltration [10], then alternative approaches should be pursued. Possible improvements of the method could come from a more targeted selection of the reference subject, which as shown by the reported results (Fig. 5) can increase the accuracy of the approach both in terms of individual muscle volume and DSC. Further studies are needed to test this hypothesis.

The geometry of the 23 muscles was captured moderately well only in the optimal combination of subjects (those with greatest lower quartile), with mean DSC of 0.74 and IQR range of 0.71 < DSC < 0.77. However, this is significantly smaller than the inter-operator dependence of the manual process, which, within the literature [2,15,21,24] is consistently found to be DSCs of around 0.90 for the muscles considered in this study. While the pair of subjects leading to the best results in terms of DSC were the most similar in terms of height and BMI, these anthropometric characteristics were very different in the pair having the second-best DSC (mean = 0.74, IQR of 0.69 < DSC < 0.79). This suggests that the newly proposed masking process (code will be made available on GitHub) achieved the goal of homogenising the subject imaging data and could be adapted for the removal of unwanted artefacts from within medical or indeed any other images.

Particularly successful approaches within the literature that used 3D deformable image registration to perform muscle segmentation were those based on longitudinal data, such as Le Troter et al. [33] and Fontana et al. [27], who attained average DSC of 0.90 and 0.85, respectively. Similar to the latter, were the DSC values here found when registering the left to right limb in the same subject. Notably, these approaches still require the manual segmentation of each subject at the baseline. Moreover, these studies segmented fewer muscles than the 23 presented in this study. Last but not least, the images collected within this study were not optimised for muscle segmentation.

Overall, the main limitation of the proposed method clearly lies in the non-satisfactory capture of individual muscle volume. These could have been caused by the propagation of inaccuracies associated with the manual segmentations of the reference images through the registration. However, this aspect is likely to be negligible since the muscles with high inter-operator variability [15] were discarded at source. More likely, the issue lied in the fact that the muscle-muscle boundaries present a weak grey-level gradient, in contrast to the muscle-fat boundaries, which are shown to have a strong grey-level gradient within the MR images (Fig. 1, 2, 4). Since ShIRT accounts for grey-level gradients within the inputted images [41], the muscle-fat and muscle-bone boundaries were registered to a higher degree of accuracy than the muscle-muscle boundaries. This unbalance in the accuracy of the registration of the different regions is highlighted by the greater RVE of the individual muscles, when compared to the total volume error. Finally, another source of error could lie within the optimisation process of the registration parameters (NS and smoothing coefficient) [41]. While in this study these parameters were optimised for the highest overall performance in segmentation accuracy across all considered lower limb muscles, the values could be optimised for the different areas of the limb. This was not implemented in this study as a rewriting of the registration toolkit would be required.

Despite the above limitations, the image registration protocol here proposed proved clearly useful when adopted to generate an augmented imaging database of 69 subjects having a much broader range of muscle volumes and geometries than the original 11 subject database. This result came after removing 41 non anatomically realistic datasets, which required some manual checking the augmented datasets, suggesting that similar care should be taken if replicating the use of the method. These datasets, made publicly available, can be used to train deep learning methods [36,37,38,39]. Machine learning and deep learning methods are now dominant tools used within the field of medical image segmentation [21,24,48]. Where the average DSC found amongst the 23 muscles considered within the present study were found to be around 0.75, considering only the optimal reference subject for each target subject, deep learning methods have been used to segment the lower limb muscles with average DSC between 0.85 [21] and 0.90 [24]. These tools are typically only suitable for studies with extremely large cohorts, but this problem has been alleviated within some medical image analysis fields, such brain tumour assessment [38] and bone segmentation [39], through data augmentation. However, this technique is yet to have been explored for muscle segmentation and the database here presented will hopefully foster efforts in this direction. To the best of our knowledge, in fact, this is the first study providing a vast, multi-operator assessed set of fully segmented, labelled augmented MR imaging sequences of the lower limb. In future work, these augmented datasets will be used to calibrate CNN models, with the potential to increase segmentation accuracy [36,37] and lead to a solution for the automatic segmentation and characterisation of muscles in vivo.

## 5. Conclusion

This study presented a novel, fully automatic muscle segmentation method using image registration, aimed at segmenting all lower limb muscles simultaneously. The results fit in well with those studies within the literature that also use image registration to segment muscle sections or individual muscles. The 3D deformable image registration is limited in its capacity to perform individual automatic muscle segmentation with a high accuracy. Nevertheless, this approach can be useful to provide total muscle volume and can be further optimised by increasing the number of reference datasets. Moreover, the publicly available augmented database built in this work would enhance any future study that would aim to use deep learning approaches for the segmentation of muscles from MR images.

## 6. Acknowledgements

The study was partially funded by Engineering and Physical Sciences Research Council (EPSRC) Frontier Multisim Grant (EP/K03877X/1 and EP/S032940/1). The authors would like to extend specific thanks to Dr. Erica Montefiori and Dr. Claude Fiifi Hayford for their assistance in this project. Lastly, we are particularly grateful for the participants that volunteered for the study.

## 8. Supporting documents

**S1 Table. Muscle volumes of the 5 subjects automatically segmented in the study**.

**S2 Appendix. Sensitivity analysis of the two registration parameters: Nodal spacing, and the smoothing coefficient**.

**S3 Appendix. Visualisation of augmented datasets for one target subject**.

**S4 Appendix. Comparisons of muscle volumes within the original and the augmented databases**.

## References

1. Pandy MG, Andriacchi TP. Muscle and joint function in human locomotion. Annual Rev Biomed Eng. 2010 Aug 15;12:401–33. doi: 10.1146/annurev-bioeng-070909-105259. PMID: 20617942.

2. Modenese L, Montefiori E, Wang A, Wesarg S, Viceconti M, Mazzà C. Investigation of the dependence of joint contact forces on musculotendon parameters using a codified workflow for image-based modelling. J Biomech. 2018 May 17;73:108–118. doi: 10.1016/j.jbiomech.2018.03.039. Epub 2018 Mar 30. PMID: 29673935

3. Hamrick MW, McGee-Lawrence ME, Frechette, DM. Fatty infiltration of skeletal muscle: mechanisms and comparisons with bone marrow adiposity. Front. Endocrinol., 20 June 2016. doi: https://doi.org/10.3389/fendo.2016.00069

4. Larsson L, Grimby G, Karlsson J. Muscle strength and speed of movement in relation to age and muscle morphology. Journal of Applied Physiology. 1979; 46(3):451–6. doi: https://doi.org/10.1152/jappl.1979.46.3.451 PMID: 438011

5. Redl C, foehler MG, Pandy MG. Sensitivity of muscle force estimates to variations in muscle–tendon properties. Human Movement Science, Volume 26, Issue 2, 2007, Pages 306-319, ISSN 0167-9457, doi: https://doi.org/10.1016/j.humov.2007.01.008.

6. Valente G, Pitto L, Testi D, Seth A, Delp SL, Stagni R, et al. (2014) Are Subject-Specific Musculoskeletal Models Robust to the Uncertainties in Parameter Identification? PLoS ONE 9(11): e112625. doi: https://doi.org/10.1371/journal.pone.0112625

7. Aoyagi Y, Shephard RJ. Aging and muscle function. Sports Med. 1992 Dec;14(6):376–96. doi: 10.2165/00007256-199214060-00005. PMID: 1470791.

8. Yoshiko A, Hioki M, Kanehira N, Shimaoka K, Koike T, Sakakibara H, et al. Threedimensional comparison of intramuscular fat content between young and old adults. BMC medical imaging. 2017; 17(1):1–8. https://doi.org/10.1186/s12880-016-0171-7 PMID: 28056868

9. Cruz-jentoft AJ, Landi F. Sarcopenia. Clin Med (Lond) 2014 Apr; 14(2): 183–186. doi: 10.7861/clinmedicine.14-2-183 PMID: 24715131

10. Gadermayr M, Disch C, Müller M, Merhof D, Gess B. A comprehensive study on automated muscle segmentation for assessing fat infiltration in neuromuscular diseases. Magn Reson Imaging. 2018 May;48:20–26. doi: 10.1016/j.mri.2017.12.014. Epub 2017 Dec 18. PMID: 29269318.

11. Lareau-Trudel E, Le Troter A, Ghattas B, Pouget J, Attarian S, Bendahan D, et al. (2015) Muscle Quantitative MR Imaging and Clustering Analysis in Patients with Facioscapulohumeral Muscular Dystrophy Type 1. PLoS ONE 10(7): e0132717. https://doi.org/10.1371/journal.pone.0132717

12. Mercuri E, Pichiecchio A, Allsop J, Messina S, Pane M, Muntoni F. Muscle MRI in inherited neuromuscular disorders: past, present, and future. J Magn Reson Imaging. 2007 Feb;25(2):433–40. doi: 10.1002/jmri.20804. PMID: 17260395.

13. Morrow JM, Sinclair CD, Fischmann A, Machado PM, Reilly MM, Yousry TA, Thornton JS, Hanna MG. MRI biomarker assessment of neuromuscular disease progression: a prospective observational cohort study. Lancet Neurol. 2016 Jan;15(1):65–77. doi: 10.1016/S1474-4422(15)00242-2. Epub 2015 Nov 6. PMID: 26549782; PMCID: PMC4672173.

14. Sookhoo S, Mackinnon I, Bushby K, Chinnery PF, Birchall D. MRI for the demonstration of subclinical muscle involvement in muscular dystrophy. Clin Radiol. 2007 Feb;62(2):160–5. doi: 10.1016/j.crad.2006.08.012. PMID: 17207699.

15. Montefiori E, Kalkman BM, Henson WH, Paggiosi MA, McCloskey EV, Mazzà C (2020) MRI-based anatomical characterisation of lower-limb muscles in older women. PLoS ONE 15(12): e0242973. https://doi.org/10.1371/journal.pone.0242973

16. Carbone V, Fluit R, Pellikaan P, van der Krogt MM, Janssen D, Damsgaard M, Vigneron L, Feilkas T, Koopman HF, Verdonschot N. TLEM 2.0 - a comprehensive musculoskeletal geometry dataset for subject-specific modeling of lower extremity. J Biomech. 2015 Mar 18;48(5):734–41. doi: 10.1016/j.jbiomech.2014.12.034. Epub 2015 Jan 8. PMID: 25627871.

17. Larsson L, Grimby G, Karlsson J. Muscle strength and speed of movement in relation to age and muscle morphology. Journal of Applied Physiology. 1979;46(3):451–6. PMID:438011

18. Suganthi GV, Sutha J, Pavarthy M, Devi CD. Pectoral muscle segmentation in mammograms. Biomed Pharmacol J 2020;13(3).

19. Handsfield GG, Meyer CH, Hart JM, Abel MF, Blemker SS. Relationships of 35 lower limb muscles to height and body mass quantified using MRI. J Biomech. 2014 Feb 7;47(3):631–8. doi: 10.1016/j.jbiomech.2013.12.002. Epub 2013 Dec 11. PMID: 24368144.

20. Ding J, Cao P, Chang HC, et al. Deep learning-based thigh muscle segmentation for reproducible fat fraction quantification using fat–water decomposition MRI. Insights Imaging 11, 128 (2020). https://doi.org/10.1186/s13244-020-00946-8

21. Zhu J, Bolsterlee B, Chow BVY, Cai C, Herbert RD, Song Y, Meijering E. Deep learning methods for automatic segmentation of lower leg muscles and bones from MRI scans of children with and without cerebral palsy. NMR Biomed. 2021 Dec;34(12):e4609. doi: 10.1002/nbm.4609. Epub 2021 Sep 21. PMID: 34545647.

22. Arnold EM, Ward SR, Lieber RL, et al. A Model of the Lower Limb for Analysis of Human Movement. Ann Biomed Eng 38, 269–279 (2010). https://doi.org/10.1007/s10439-009-9852-5

23. Modenese L, Kohout J. Automated Generation of Three-Dimensional Complex Muscle Geometries for Use in Personalised Musculoskeletal Models. Ann Biomed Eng 48, 1793–1804 (2020). https://doi.org/10.1007/s10439-020-02490-4

24. Ni R, Meyer CH, Blemker SS, Hart JM, Feng X. Automatic segmentation of all lower limb muscles from high-resolution magnetic resonance imaging using a cascaded three-dimensional deep convolutional neural network. J Med Imaging (Bellingham). 2019 Oct;6(4):044009. doi: 10.1117/1.JMI.6.4.044009. Epub 2019 Dec 28. PMID: 31903406; PMCID: PMC6935014.

25. Engstrom CM, Fripp J, Jurcak V, Walker DG, Salvado O, Crozier S. Segmentation of the quadratus lumborum muscle using statistical shape modeling. J Magn Reson Imaging. 2011 Jun;33(6):1422–9. doi: 10.1002/jmri.22188. PMID: 21591012.

26. Engstrom CM, Walker DG, Kippers V, Mehnert AJ. Quadratus lumborum asymmetry and L4 pars injury in fast bowlers: a prospective MR study. Med Sci Sports Exerc. 2007 Jun;39(6):910–7. doi: 10.1249/mss.0b013e3180408e25. PMID: 17545879.

27. Fontana L, Mastropietro A, Scalco E, et al. Multi-Steps Registration Protocol for Multimodal MR Images of Hip Skeletal Muscles in a Longitudinal Study. Appl. Sci. 2020, 10(21), 7823; https://doi.org/10.3390/app10217823

28. Hides J, Stanton W, Freke M, Wilson S, McMahon S, Richardson CA. MRI study of the size, symmetry and function of the trunk muscles among elite cricketers with and without low back pain. Br J Sports Med 2008; 42: 809–813.

29. Lexell J, Taylor CC. Variability in muscle fibre areas in whole human quadriceps muscle: effects of increasing age. J Anat. 1991 Feb;174:239-49. PMID: 2032938; PMCID: PMC1256058.

30. Duda GN, Brand D, Freitag S, Lierse W, Schneider E. Variability of femoral muscle attachments. J Biomech. 1996 Sep;29(9):1185–90. doi: 10.1016/0021-9290(96)00025-5. PMID: 8872275.

31. Holzbaur KR, Murray WM, Gold GE, Delp SL. Upper limb muscle volumes in adult subjects. J Biomech. 2007;40(4):742–9. doi: 10.1016/j.jbiomech.2006.11.011. Epub 2007 Jan 22. PMID: 17241636.

32. Ogier A, Sdika M, Foure A, Le Troter A, Bendahan D. Individual muscle segmentation in MR images: A 3D propagation through 2D non-linear registration approaches. Annu Int Conf IEEE Eng Med Biol Soc. 2017 Jul;2017:317–320. doi: 10.1109/EMBC.2017.8036826. PMID: 29059874.

33. Le Troter A, Fouré A, Guye M, Confort-Gouny S, Mattei JP, Gondin J, Salort-Campana E, Bendahan D. Volume measurements of individual muscles in human quadriceps femoris using atlas-based segmentation approaches. MAGMA. 2016 Apr;29(2):245–57. doi: 10.1007/s10334-016-0535-6. Epub 2016 Mar 16. PMID: 26983429.

34. Fontana, L, Mastropietro A, Scalco E, et al. Multi-Steps Registration Protocol for Multi-modal MR Images of Hip Skeletal Muscles in a Longitudinal Study. November 2020, Applied Sciences 10(21):7823 doi: 10.3390/app10217823

35. Hesamian MH, Jia W, He X, et al. Deep Learning Techniques for Medical Image Segmentation: Achievements and Challenges. J Digit Imaging 32, 582–596 (2019). https://doi.org/10.1007/s10278-019-00227-x]

36. Shen Z, Xu Z, Olut S, Niethammer M. Anatomical Data Augmentation via Fluid-based Image Registration. arXiv 2020 doi: 10.48550/ARXIV.2007.02447

37. Shorten C, Khoshgoftaar TM. A survey on Image Data Augmentation for Deep Learning. J Big Data 6, 60 (2019). https://doi.org/10.1186/s40537-019-0197-0

38. Nalepa J, Marcinkiewicz M, Kawulok M. Data augmentation for brain-tumor segmentation: a review. Front. Comput. Neurosci., December 2019 doi: https://doi.org/10.3389/fncom.2019.00083

39. Noguchi S, Nishio M, Yakami M, Nakagomi K, Togashi K. Bone segmentation on whole-body CT using convolutional neural network with novel data augmentation techniques. Computers in Biology and Medicine, Volume 121, 2020, 103767, ISSN 0010-4825, doi: https://doi.org/10.1016/j.compbiomed.2020.103767.

40. Canny J. A Computational Approach To Edge Detection, IEEE Transactions on Pattern Analysis and Machine Intelligence, 8(6):679–698, 1986.

41. Barber DC, Hose DR. Automatic segmentation of medical images using image registration: diagnostic and simulation applications. J Med Eng Technol. 2005 Mar-Apr;29(2):53–63. doi: 10.1080/03091900412331289889. PMID: 15804853.

42. Barber DC, Oubel E, Frangi AF, Hose DR. Efficient computational fluid dynamics mesh generation by image registration. Medical Image Analysis, Volume 11, Issue 6, 2007, Pages 648–662, ISSN 1361-8415, doi: https://doi.org/10.1016/j.media.2007.06.011.

43. Dall’Ara E, Barber D, Viceconti M, About the inevitable compromise between spatial resolution and accuracy of strain measurement for bone tissue: A 3D zero-strain study,Journal of Biomechanics,Volume 47, Issue 12, 2014, Pages 2956-2963, ISSN 0021-9290,https://doi.org/10.1016/j.jbiomech.2014.07.019.

44. Hayford CF, Montefiori E, Pratt E, Mazzà C. Predicting longitudinal changes in joint contact forces in a juvenile population: scaled generic versus subject-specific musculoskeletal models. Comput Methods Biomech Biomed Engin. 2020 Oct;23(13):1014–1025. doi: 10.1080/10255842.2020.1783659. Epub 2020 Jun 26. PMID: 32588655.

45. Dice LR. (1945). Measures of the Amount of Ecologic Association Between Species. Ecology. 26 (3): 297–302. doi:10.2307/1932409. JSTOR 1932409.

46. Rockafellar R, Wets I, Roger JB, (2005). Variational Analysis. Springer-Verlag. p. 117. ISBN 3-540-62772-3.

47. Thelen DG. (February 14, 2003). Adjustment of Muscle Mechanics Model Parameters to Simulate Dynamic Contractions in Older Adults. ASME. J Biomech Eng. February 2003; 125(1): 70–77. doi: https://doi.org/10.1115/1.1531112

48. Lenchik L, Heacock L, Weaver AA, et al. Automated Segmentation of Tissues Using CT and MRI: A Systematic Review, Academic Radiology, Volume 26, Issue 12, 2019, Pages 1695–1706, ISSN 1076-6332, doi: https://doi.org/10.1016/j.acra.2019.07.006.

